# *Ostreococcus tauri* is a high-lipid content green algae that extrudes clustered lipid droplets

**DOI:** 10.1101/249052

**Authors:** Chuck R. Smallwood, William Chrisler, Jian-Hua Chen, Emma Patello, Mathew Thomas, Rosanne Boudreau, Axel Ekman, Hongfei Wang, Gerry McDermott, James E. Evans

## Abstract

Lipid droplet biogenesis, accumulation and secretion is an important field of research spanning biofuel feedstock production in algae and yeast to plant-microbe symbiosis or human metabolic disorders and other diseases. Here we evaluate the critical elements that influence lipid accumulation in the highly simplified and smallest known eukaryote *Ostreococcus tauri* and identify several conditions that satisfy its classification as an oleaginous green alga. In addition, these experiments revealed the release of excess lipids in pea-pod like structures where many dense lipid droplets are clustered in a linear fashion surrounded by an enveloping membrane which contrasts with known mechanisms from other eukaryotes. These results highlight the potential for *Ostreococcus tauri* to probe the evolution of lipid droplet dynamics as an emerging model organism with a compacted eukaryotic genome and also to impact lipid feedstock bioproduction applications either directly or using synthetic biology.

**One Sentence Summary:** The smallest known eukaryote *Ostreococcus tauri* is oleaginous and sheds lipid droplets as pea-pod like membrane enclosed clusters.

## Main Text

Bioproducts from oleaginous green algae include food additives, cosmetics, commodity chemicals and lipid droplet feedstocks for biofuels ^1^. Recent reviews of bioprocessing described the costs associated with biomass production as the primary bottleneck for achieving competitive pricing while low-to no-energy harvesting options can also improve the cost-benefit by efficiently isolating the bioproduct from cellular material ^2,3^. To accelerate more cost-effective bioproduction of lipid feedstocks it is imperative to identify new species or cellular mechanisms that can be exploited to enhance production or harvesting. Lipid droplets and their dynamics are also important for a range of human health related processes with roles in lipid storage, metabolism, signaling, inflammation and cancer ^4^.

*Ostreococcus tauri* is phototrophic and the smallest known free-living eukaryote ^5^. Found in most oceans worldwide, this dominant picoplankton can thrive in adverse environments such as copper contaminated seawater ^6^. Under standard growth conditions, the cell dimensions are around 0.8-1.0 micron in thickness and about 1.0-1.4 microns in diameter and this houses a highly simplified ultrastructure with a single nucleus, chloroplast and mitochondrion while lacking both a cell wall and flagella ^7^. *O. tauri* also has a highly compacted genome with just under 8,000 genes ^8,9^. The simplified state of both the cell architecture and its genome, combined with published protocols for genetic tractability have positioned *O. tauri* as an emerging model organism that has already informed mechanistic details underlying circadian rhythm, cell division, and modeling kinome networks ^10-13^. We posited that the compact nature of *O. tauri* could also reveal new insights to lipid droplet dynamics.

In many eukaryotes, lipid accumulation is inducible by modulating light intensity, altering temperature, pH, salinity or nutrient availability ^14-16^. For example, different algae species have been shown to trigger neutral lipid (NL) accumulation upon iron, phosphorous, silicon (specifically for diatoms), sulfur and nitrogen nutrient starvation ^17,21^. However, like many biological processes that evolved across diverse microorganisms, the precise mechanisms underlying NL accumulation (specifically triacylglycerols) due to nutrient availability varies widely and in many cases, remains obscure ^15,22,23^. Interestingly, the standard growth media for *O. tauri*, Keller (K) media, includes several nutrients (N, P, S, Fe) at concentrations around 10^-4^–10^-5^ M that are known to induce triacylglycerol accumulation in other algae and oleaginous eukaryotes. Co-factors (Mo, Co, Mn, Zn) and vitamins (B1, B7, B12) are also present in K media but at much lower concentration around 10^-7^–10^-9^M ^24^. Prior studies comparing growth of *O. tauri* under normal conditions versus nitrogen (N) starvation at 10% of normal nitrogen availability (media containing 10^-5^ instead of 10^-4^M combined NH_4_ & NO_3_) have reported some elevated NL accumulation providing support that *O. tauri* is susceptible to similar environmental cues ^23,25,26^. The presence of sulfur (S), phosphorus (P), and iron (Fe) in K-media at similar concentration as nitrogen therefore elicited the question of whether these other nutrients are truly required at elevated levels for normal growth of *O. tauri* and if their single or multiplexed removal would enhance lipid droplet production beyond simple N starvation.

For simplicity, we primarily focused this study on evaluating the effects of each major component in *O. tauri’s* standard growth media as nutrient starvation has typically yielded the highest levels of triacylglycerol feedstocks in other microalgae, diatoms and fungi. The simultaneous and complete depletion of both forms of available N (ammonium and nitrate) present in K media, induced NL production (Fig. 1) similar to previous reports ^14,27^. Interestingly, media containing only nitrate as the bioavailable N source grew normal to the standard K media and had elevated but diffuse lipid signal. Whereas media containing only ammonium resulted in discrete lipid droplet formation, but to a lesser extent compared to total N deficient media. Phosphorous deprivation also triggered detectable NL accumulation while removal of sulfur and iron from Keller media exhibited minimal to no increases in lipid production and showed relatively normal growth (Fig. 1). As reported in other organisms, lowered total biomass yields were observed for simple N and P depletion.

**Fig. 1:**
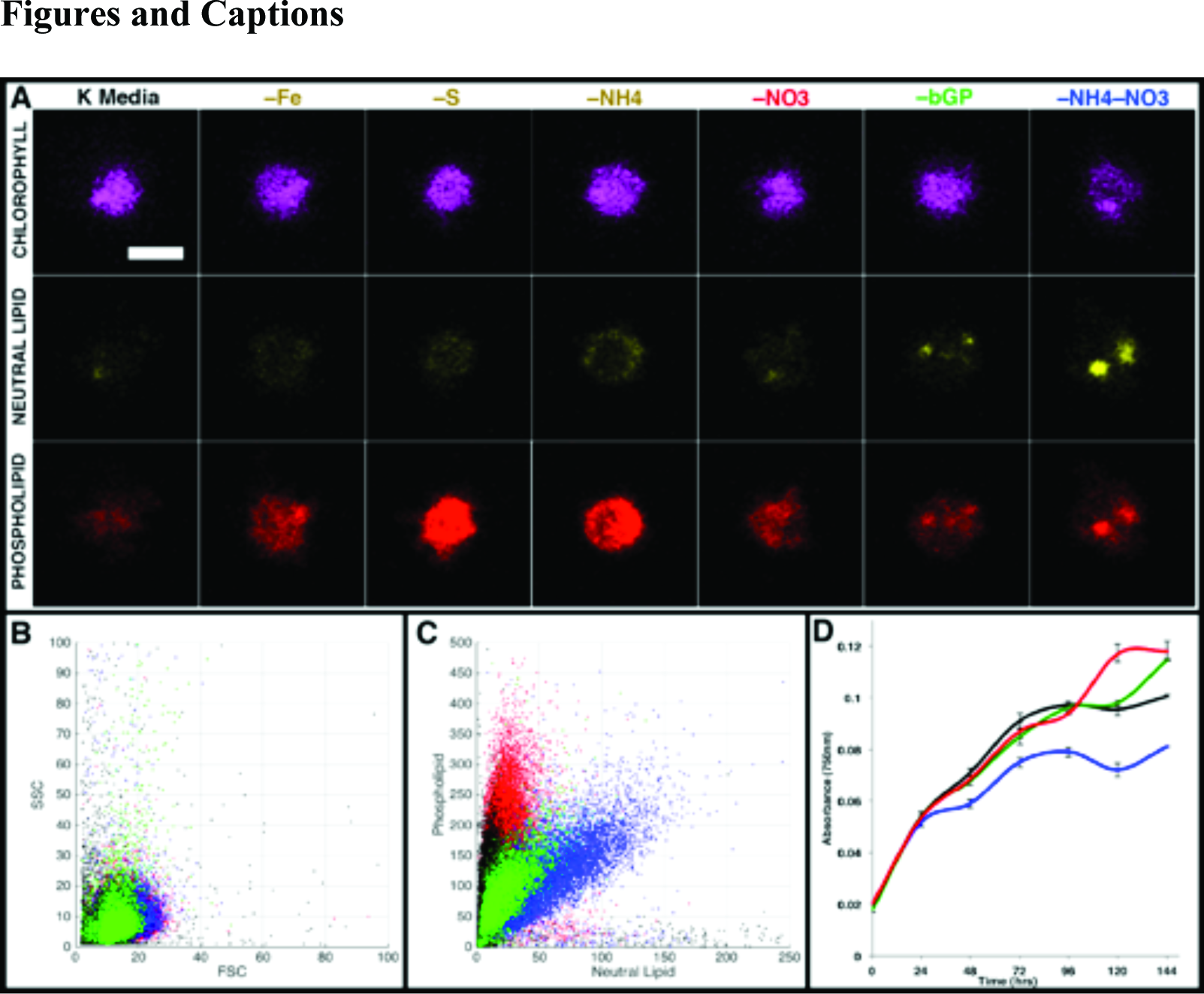
Nutrient deprivation of phosphate and nitrogen induces lipid droplet formation. Confocal fluorescence microscopy (1μm scale bar) of *O. tauri* cells stained with Nile Red after 72 hours of culturing in defined media conditions. B) Cell culture growth monitored at 750nm with colors corresponding to the labels in (A). Same cell cultures were analyzed by FACS to obtain side-scattered light (SSC) vs forward-scattered light (FSC) (C) and PL versus NL (D) plots.

Since complete N starvation showed elevated levels of NL accumulation, we evaluated the impact of variable N availability. Discrete lipid droplets were first detected in cells exposed to media lacking 80% of normal total N levels (Supp. Fig. 1). Further decreasing N availability led to less total biomass and lowered chlorophyll autofluorescence but increased the size and number of detected lipid droplets as well as cell size. Based on a trade-off between biomass and lipid yields, removal of 90% and 100% of normal N concentration were identified as favorable baselines for single nutrient starvation conditions. Given that complete removal of P from the media also showed elevated NL accumulation, albeit significantly less than simple N deprivation, we explored whether the simultaneous depletion of both N and P could increase LD feedstocks further. Variable levels of P starvation using K media deprived of 90% N as the base were compared (Sup. Fig. 2) and discrete LDs were first detected in cells grown in media lacking 60% of normal P levels. LD content increased with more severe starvation.

Although the increased lipid signal upon concurrent N and P starvation was intriguing, we also considered that the removal of the sole P present in the K media (beta-glycerol phosphate (bGP)) could also potentially deplete a bioactive form of glycerol. While *O. tauri* was expected to be a photoautotroph, other microalgae have been found to grow on glycerol as a carbon source ^28^. Per its annotated genome, *O. tauri* lacks any pathway for direct utilization of glycerol-2-phosphate or annotated genes for bGP uptake. However, we discovered the presence of an alkaline phosphatase gene (0t10g02060) in the KEGG database that likely participates in the folate biosynthesis pathway ^29^. We hypothesized that *O. tauri* could potentially utilize the same alkaline phosphatase to enzymatically cleave phosphate from bGP extracellularly and subsequently generate pools of bioavailable P and glycerol that could each be taken up by the cell. To test this, we supplemented increasing levels of glycerol into our cell cultures while simultaneous depleting bGP in the K-90%N and K-100%N media. We found that increasing the glycerol concentration up to 50mM (well beyond that of normal bGP concentrations), resulted in significant increases of intracellular lipids while showing elevated (not depressed) biomass yields (Fig. 2 & Supp. Fig. 3). Beyond 50mM glycerol slower biomass accumulation is seen which likely results from detrimental osmotic effects impacting cell viability. While the bioimaging data clearly showed a dramatic increase in total lipid content, we also quantified the best lipid production condition from this screen. We subjected the cultures grown in K-100%N, K-100%N-100%P+50mMglycerol, K-90%M–60%bGP+50mMglycerol and K-90%M–60%bGP+20mMglycerol to conventional lipid extraction protocols. These conditions yielded total lipid content (dry lipid weight / dry biomass weight) of 24.7%, 25.7%, 31.2% and 30.8% respectively. Although the total lipid content appears similar for these conditions, it should be noted that the dry biomass total weight for the K-90%M–60%bGP+50mMglycerol was 146% that of K-100%N. The measured lipid content for both the K-90%M–60%bGP+50mMglycerol and K-90%M–60%bGP+20mMglycerol conditions surpass the 30% threshold for classification as a high-lipid content or oleaginous organism after only 4 days of starvation.

**Fig. 2:**
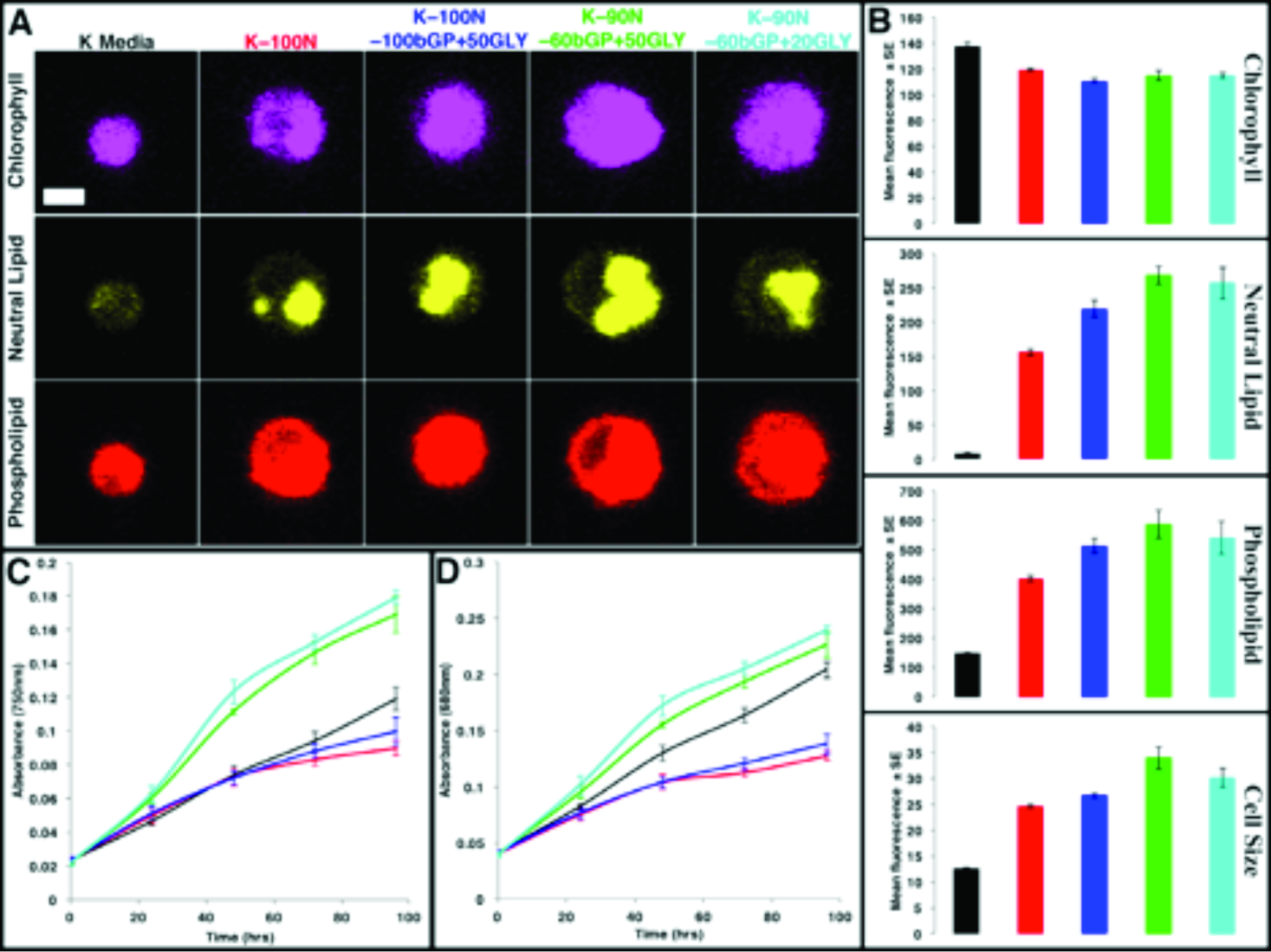
Conditions for high biomass and neutral lipid accumulation. A) Confocal fluorescence microscopy (1μm scale) of *O. tauri* cells stained with Nile red after 96 hours of culturing in defined K media conditions. B) The same cell cultures were analyzed by FACS to obtain population level data. Cell culture growth was monitored for chlorophyll content at 680nm (C) and particulates at 750nm (D). All colors representative of labels from (A).

**Fig. 3:**
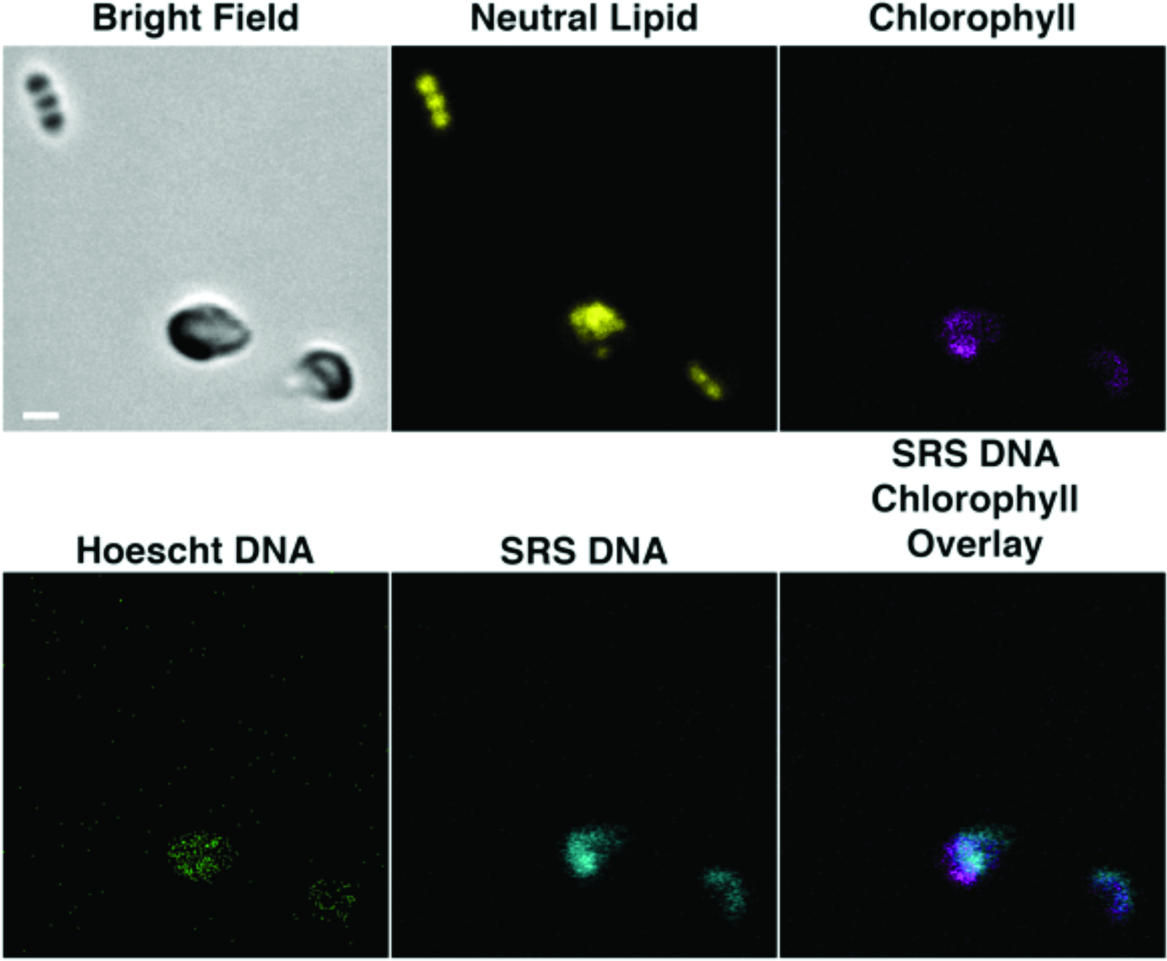
Confirmation of no nucleic acid or chlorophyll autofluorescence in the pea-pod extruded structures. The bright field image depicts healthy *O. tauri* cells (grown in K-90%N-60%P+20mMGlycerol) in the and pea-pod like extrusions. No detectable signal for DNA or chlorophyll was detected in the LD containing pea-pod structures using both label-based and label-free imaging approaches.

Surprisingly, bioimaging experiments also uncovered a previously unreported mechanism for lipid droplet extrusion. Eukaryotes typically release individual LDs if they release/secrete the LDs at all ^30,31^. Initially, confocal experiments described above revealed conditions in which “pea-pod” like structures were present in the culture. These structures stained positive for NL and phospholipid but did not possess any chlorophyll autofluorescence (Fig. 3). Staining of nucleic acids with Hoechst labeling also failed to label nucleic acid in these pea-pod structures but did stain the nucleus of normal *O. tauri* cells. Similarly, the use of label-free Stimulated Raman Scattering microscopy on the same cells (tuned to a common nucleic acid peak at 1093 cm^-1^) again failed to identify any nucleic acid signatures in the pea-pod structures (Fig. 3).

A few of the confocal image sets showed evidence of oriented lipid droplets in blebs still fused to the main *O. tauri* cells (Fig. 4a). Thus, it seemed plausible that the free-floating pea-pod structures originated from *O. tauri*, and were probably formed through a concerted cell blebbing and unknown extrusion mechanism where clustered lipid droplets were released into solution surrounded by an enclosing outer phospholipid membrane. This sloughing mechanism does not appear related to cell death as longer incubation periods show continued biomass increases and stable chlorophyll per cell ratios. Additionally, although a variable lag time is present, all cultures tested here showed the ability to recover normal growth following a batch dilution into normal K media (Supplemental Figure 4).

**Fig. 4:**
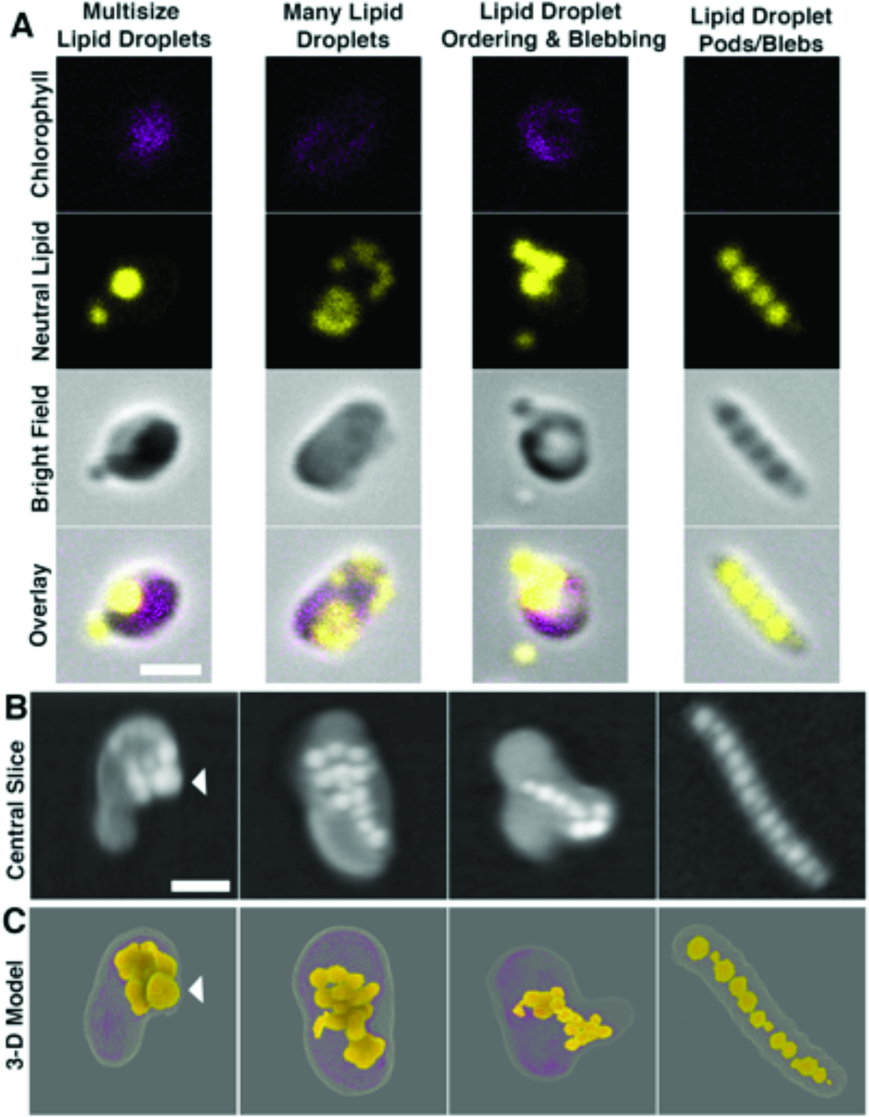
Visualizing various cellular ultrastructures and lipid droplet arrangements. Confocal microscopy (A) of K-90%N-60%bGP+20mMGlycerol cultures at 96 hours were stained with Nile Red to reveal *O. tauri* cells containing various distributions of lipids such as multisize lipid droplets, many lipid droplets, ordered lipid droplets with lipid droplets outside the cell, and pea pod-like lipid structures. Central slices from the nanotomography reconstructions (B) and segmented 3-D models (C) highlight examples of the various lipid droplet sizes and distributions that included multisize lipid droplets, ordered lipid droplets extrusion events and pea pod-like structures similar in organization and appearance to those in (A). White bars represent 1 μm scale and white arrow identifies same large lipid body from central slice to segmented 3-D model.

Thus, to independently confirm the detection of LDs in pea-pod like structures we performed cryogenic soft X-ray nanotomography which is a quantitative imaging method that can compare the density of lipid accumulations under different environmental conditions at the single cell level. We indeed visualized free-floating pea-pod like structures containing only densities with linear absorption coefficients matching lipid droplets and even captured several cells wherein the intracellular lipid droplets were organized in a linear arrangement prior to their partitioning into a bleb-like structure (Fig. 4b and Sup. Movie 1).

Although the standard K media contains similar concentrations of iron, phosphorus, sulfur, nitrate and ammonium, only depletion of N and P stimulated lipid droplet production for *O. tauri.* While the detailed molecular mechanisms regulating lipid accumulation and oriented extrusion continue to remain elusive for this organism, these results highlight a convenient route for induction of high lipid content with concomitant high biomass yields in *O. tauri.* This has not yet been reported in other microalgae or yeast where there is typically a trade-off between high-lipid content and biomass accumulation. Interestingly, an NCBI BLAST search of *O. tauri* against oleosin, perilipin, and major lipid droplet proteins typically associated with the surface of lipid droplets in other algae and higher eukaryotes shows no significant homologs. This lack of canonical genes associated with the surface of lipid droplets or their secretion as well as direct evidence of the pea-pod like extrusion suggests that this ancient organism may be employing a novel mechanism for offloading excess lipids when starved of both N and P and supplemented with carbon.

This organism or components of its pathway may therefore have direct applications towards enhancing industrial bioproduction of lipid feedstocks. Furthermore, the discovery of the extrusion of pea-pod like structures containing only clustered lipid droplets makes this organism an intriguing candidate model system for understanding eukaryotic lipid dynamics especially due to its simplified ultrastructure and genome. For example, future work can exploit the highly reduced pathways present in *Ostreococcus* and its genetic tractability to interrogate how the deletion of specific endogenous genes or addition of exogenous genes may force the phenotype of pea-pod like lipid droplet extrusion to more closely resemble that of higher eukaryote lipid secretion to better disentangle the complexities of lipid droplet dynamics central to processes impacting energy, health and the environment.

## Materials and Methods

### Media, strain maintenance, growth and starvation conditions

*O. tauri* cell cultures were obtained from the Roscoff Culture Collection (RCC745); strain name: 0TTH0595, which has been fully sequenced ^8^. Cultures of RCC745 were cultured in Keller media or respective K media with or without specified nutrients to create defined media conditions ^24^. All K media based culture conditions were prepared in fresh artificial seawater (ASW) with defined amounts of nutrients. Our defined media conditions include nitrogen and elemental starvation and in some cases supplementation with excess carbon in the form of glycerol. The defined conditions for nitrogen limitation are straight forward, using the base K media by limiting nitrogen sources and supplementing with percent molar amounts of both ammonium and nitrate to create a gradient survey (–100, –90, –80, –60, –20 and –0%) of limiting nitrogen, whereas our –0% has no nitrogen removed from normal K media and –100% has no ammonium or nitrate sources present. Defined conditions for elemental starvation was carried out by first growing the cells to mid to late log phase in normal K media then gently centrifuging cultures, washing and suspending them into defined K media that was prepared without specific sources of the following ‒Fe, ‒S, ‒Mg, ‒*β* ‒glycerophosphate (-BGP), ‒NO3, ‒NH4, ‒NO3-NH4, and ‒BGP+Glycerol. We recorded and displayed absorbance at 680nm and 750nm to obtain a measure of both chlorophyll content and particulate matter, respectively, for each culture. Graphing the ratio of 750nm/680nm provided a measure of chlorophyll functional efficiency as well as possible lipid metabolic flux. Graphical analysis of each growth and starvation curve required minimal normalization due to our consistent efforts in capturing cells during mid-log stages of growth. For each experiment replicate *O. tauri* cultures were grown up to mid to late log phase under blue diurnal light (18–20 μmol photons/m^2^/s) in Percival Scientific light incubator (Iowa, USA), captured before stationary phase, to ensure fresh healthy cells prior to nutrient limitation, because if allowed to enter stationary phase some lipid accumulation can be observed. To prepare cell cultures for starvation survey we gently centrifuged fresh cultures at 2200xg for 10 mins with swing bucket rotor centrifugation and washed cell pellets once with defined K media of interest, then suspended cells in defined media conditions and continued diurnal light entrainment for specified time courses in sealable CytoOne non-treated cell culture flasks (USA Scientific, USA) with mixing of cultures once per 24 hours.

### Fluorescence activated cell sorting analysis of intracellular lipid content

*O. tauri* cells were cultured to mid-log phase and gently centrifuged to concentration then stained with Nile Red for exactly 10 mins before each experimental measurement on the BD INFLUX flow cytometer (BD Biosciences, San Jose, CA, USA). Forward Scatter (FSC) and Side Scatter (SSC) were used to gate out any non-specific cellular debris. We analyzed specific gating in the range of known cell size of *O. tauri* to determine the fluorescence from stained neutral lipid (488/542±13.5 nm), phospholipid (561/615±12 nm), and natural chlorophyll autofluorescence (640/670±15 nm) for defined populations of cells. Each individual FACS experiment was calibrated to 3.6 side scatter 10 mins before running our sample measurements in defined media cultures. We plotted the fluorescence intensity of neutral lipid and phospholipid fluorescence intensity at specific time points in scatter plots to demonstrate population dynamics for each sample condition at 72 hours into starvation. We used standard K media as a control for normal conditions and nominal lipid staining for comparison to high lipid content conditions.

### Fluorescence and SRS confocal microscopy

Confocal images were obtained on Zeiss LSM 710 (Carl Zeiss AG, Germany) confocal microscope with a 100x oil immersion objective. An InTune Laser with 505nm and 535nm light was used to maximize the separation of the triglyceride (585nm) and phospholipid (638nm) emission peaks while diminishing crosstalk of the Nile Red Stained cells. In addition, we excited chlorophyll autofluorescence with 405nm light and monitored the emission profile at 680nm. Nile Red stained cells were immobilized on glass slides with poly-L-lysine and imaged immediately with z-scan slicing of 0.43 μm to survey whole cell fluorescence labeling distribution. All fluorescence channels were set with identical gain and laser power settings to provide relative levels of fluorescent intensity. We have displayed selected slices from our z-scans and applied no adjustments of contrast or gain during post processing. Microalgae cells were stained with Hoescht 33342 DNA and Nile Red stains for imaging on a Leica SP8 Confocal Microscope using a 63× 1.40 NA water immersion objective with 1 × and 4× zoom to satisfy Nyquist frequency. Auto-fluorescence of chlorophyll (ex: 552 nm, Em: 675-690nm) and fluorescence of Hoescht (Ex: 405, Em: 420-520nm), neutral lipid (Ex: 488, Em: 570-600nm) and phospholipid (Ex: 488, Em: 630-647nm) were monitored with bright-field channels during sequential scans with Leica HyD photon detector. SRS confocal images were obtained using a 63x 1.40na objective with excitation from an APE Pico Emerald laser source with 6 picosecond pulse widths, 1032 fundamental laser, and OPO at 926.6nm (SRS DNA) with dual 850+/-10nm short pass filters to visualize intracellular nucleic acid content false colored in cyan (Fig 3. bottom panel).

### Soft X-ray nanotomography, reconstruction and segmentation

*O. tauri* cultures were grown up in standard K media to mid-log phase under blue diurnal light (18–20 μmol photons/m^2^/s). To prepare cell cultures for starvation survey we gently centrifuged fresh cultures at 2200xg for 10 mins with swing bucket rotor centrifugation and washed cell pellets once with defined K media of interest then suspended cells in defined media conditions and continued diurnal light entrainment for specified time courses in sealed tubes for transport to Advanced Light Source facilities. Thin-walled glass capillaries to hold the cells for 3D imaging were pulled and assembled as described previously ^32^. Soft x-ray tomographic imaging was carried out on cryopreserved specimens to mitigate the effects of radiation damage and prevent movement of fine structural details during data acquisition ^33,34^. Prior to imaging, cells were pelleted to a high titer and loaded into tapered specimen capillaries (~5-6 μm in diameter at the tip) using a standard micropipette. Once loaded into capillaries, cells were immediately cryopreserved by rapid plunging into liquid propane using a custom fast-freezing apparatus.

Cells were mixed with 6μm beads to maintain cellular integrity during crypreservation. Frozen specimens were cryo-transferred into custom storage boxes using a home-built cryo-transfer device and stored in liquid nitrogen.

Soft x-ray data acquisition was carried out on beamline 2.1, a soft x-ray microscope in the National Center for X-Ray Tomography (NCXT) located at the Advanced Light Source in Berkeley, California ^35^. The microscope soft x-ray illumination was generated by a bend-magnet in the synchrotron lattice and focused onto the specimen by a Fresnel Zone Plate (FZP) condenser. Specimen illumination was order-sorted by a pinhole positioned just in front of the specimen. A second zone plate, located downstream of the specimen, magnified and focused an image of the specimen on a CCD detector. During data collection, the cells were maintained in a stream of helium gas that had been cooled to liquid nitrogen temperatures. Each tomographic dataset (i.e., 90 projection images spanning a range of 180°) was collected using Fresnel zone plate based objective lens with a resolution of 50 nm. Exposure times for each projection image ranged from 150 to 300 msec. The software suite AREC3D ^36^ was used to align the projection images calculate tomographic reconstructions.

Tomography segmentation was completed using in-house code written in MATLAB (MATLAB 9.1, The MathWorks Inc., Natick, MA, 2017). The segmentation primarily relies on threshold ranges for different intracellular structures of interest. The segmenation code generates montages for each cell membrane, lipid bodies and chloroplast which are then used to generate the 3-D models and animations in Drishti ^37^. For the supplemental video, the montages generated by the MATLAB code for the cell, chloroplast and lipid bodies were overlaid using the MIJ library that provides the platform to integrate MATLAB and ImageJ ^38^. The 2-D animations were generated using ImageJ and 3-D animations using Drishti. Slices of the 3-D animation were embedded to the video using ImageJ.

### Quantification of Cell Biomass and Small Batch Lipid Extraction

Cell cultures were grown and resuspended into starvation and/or supplementation conditions and allowed to incubate for 72 hours before cell density measurements and harvesting. We harvested and partially dried whole culture content using 0.45 micron PVDF filters (Merck Millipore, USA) with a vacuum pressure at 25 psi. We dried the cell biomass before weighing on an analytical balance by using a turbo pump to pull a vacuum down to 25 inHg. The dry weight of each filter was obtained before and after filtration of cell biomass to measure total cell dry weights of each culture. We placed each dried filter with cells and material attached into glass tubes for lipid extraction. The lipid extraction followed a modified procedure prescribed by Olmstead et al. to quantify total lipids in microalgal cells with the modification of 1M NaCl in the place of water ^39^. We carefully collected the bottom chloroform organic layer during each extraction containing the total lipid content and combined these extracts into pre-weighed glass tube for drying under nitrogen gas at 40°C using the Techne Sample Concentrator with model DB-3D Dri-Block (Bibby Scientific Limited, UK). Post drying, we weighed each tube for total dry lipid weight and combined the dry cell weights to determine the percentage of dry lipid content (mg) per dry cell weight (mg).

## Acknowledgements

Work was supported by DOE-BER Mesoscale to Molecules Bioimaging Project FWP# 66382. A portion of the research was performed using the Environmental Molecular Sciences Laboratory (EMSL), a national scientific user facility sponsored by the Department of Energy’s Office of Biological and Environmental Research and located at PNNL. This research also used resources of the Advanced Light Source, which is a DOE Office of Science User Facility under contract no. DE-AC02-05CH11231.

## Author contributions

C.S. and J.E. devised experiments for the study. C.S. conducted experiments. W.C., J.C., E.P., M.T., R.B., A.E., H.W., and G.M. all contributed to the study.

## Supplementary Materials

Supplemental Figures 1-4, Supplemental Movie 1

**Supplemental Figure 1:**
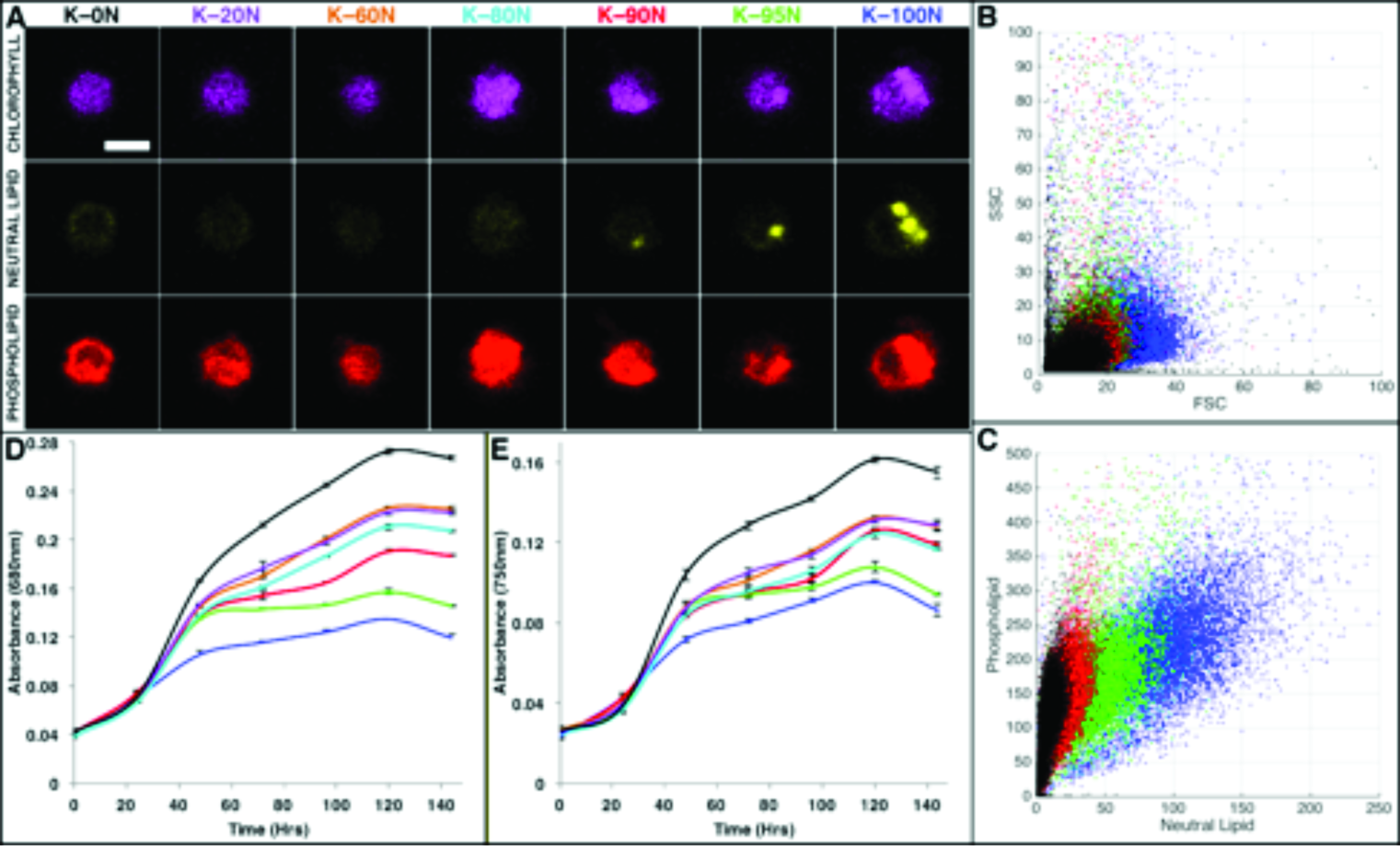
Nitrogen depletion revealed incremental increases in neutral lipid. Confocal fluorescence microscopy (A) of varying total nitrogen cell cultures stained with Nile Red revealed incremental lipid droplet formation with decreasing nitrogen content. Growth curves of cultures were collected by monitoring absorbance at 680nm (D) and 750nm (E) for chlorophyll and biomass, respectively. FACS of cultures stained with Nile Red displayed slight increases in cell size (FSC) from SSC vs FSC scatter plot (B) and increasing neutral lipid and phospholipid content with decreasing total nitrogen from the scatter plot of Phospholipid vs Neutral Lipid (C) signal.

**Supplemental Figure 2:**
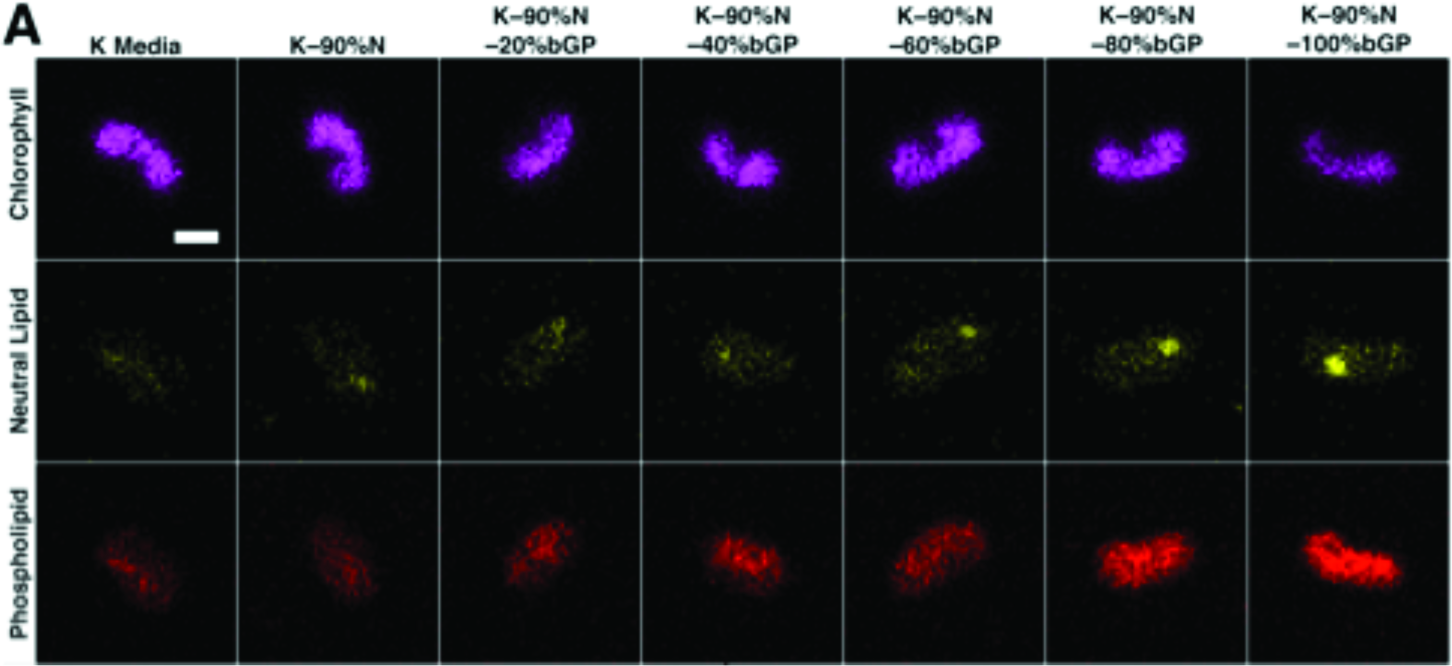
Variable phosphate depletion series with simultaneous 90% nitrogen starvation. Confocal fluorescence microscopy (A) of cultures with 90% total nitrogen depletion and incremental depletion of beta-glycerol phosphate revealed increasing neutral lipid and phospholipid content with slight decreases in chlorophyll auto-fluorescence. Cell culture growth was monitored at absorbance wavelengths 680nm (B) and 750nm (C) and displayed that beyond K-90%N–60%bGP cells suffered dramatic biomass depletion.

**Supplemental Figure 3:**
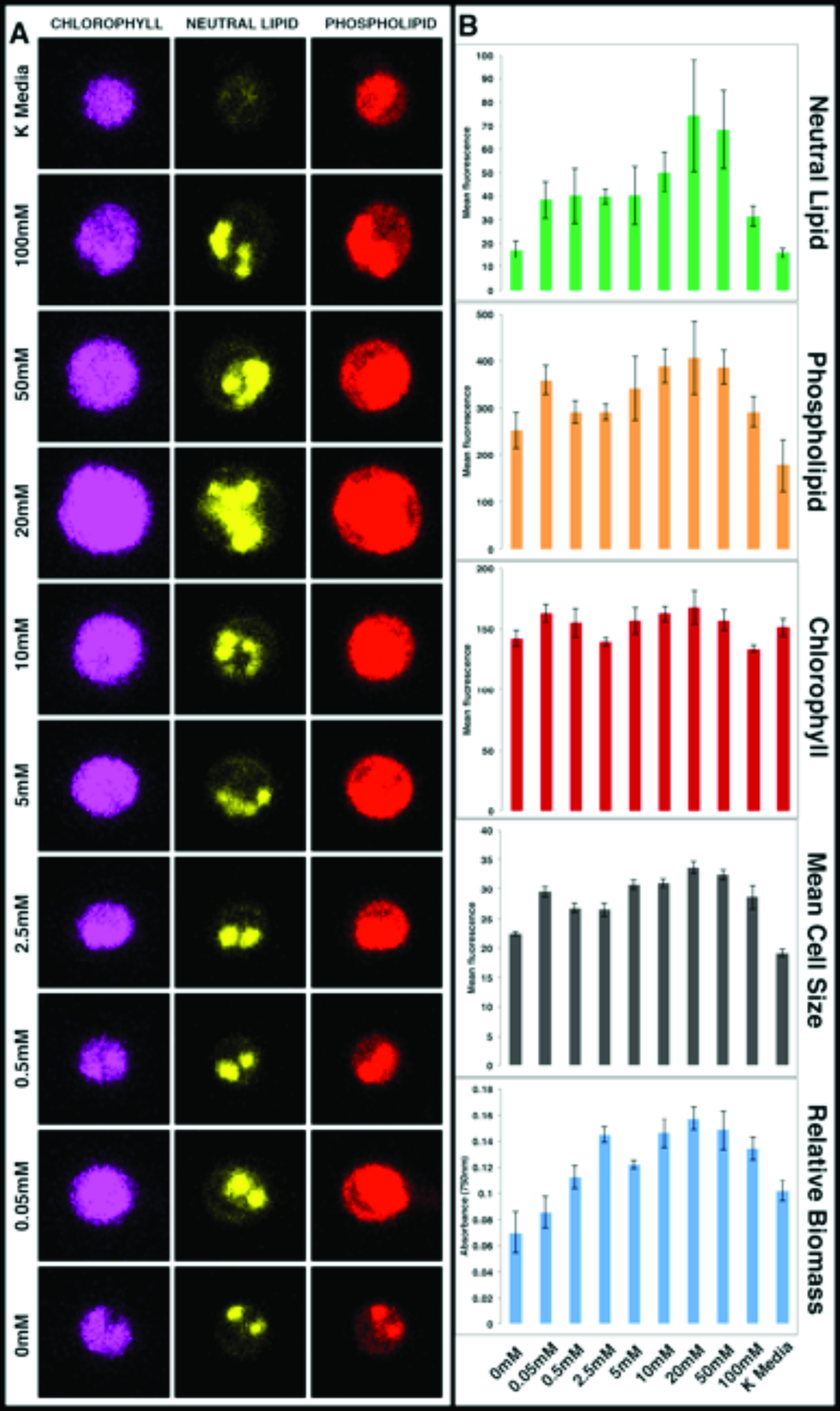
Glycerol concentration dependent cell size and TAG lipid accumulation. A) Confocal fluorescence microscopy (1μm scale bar) of *O. tauri* cells after 72 hours of culturing in defined K media condition K-90%N–100%bGP with varying glycerol concentration. B) Cell culture growth was monitored at 750nm (full data not shown) for relative biomass at 96 hours. The same cell cultures were analyzed by FACS to obtain neutral lipid (green bar graph), phospholipid fluorescence (orange bar graph), chlorophyll autofluorescence (red bar graph), forward scattered light (dark grey bar graph), and 750nm Absorbance at 72 hours (light blue bar graph). Error bars represent the error in five biological replicate cultures.

**Supplemental Figure 4:**
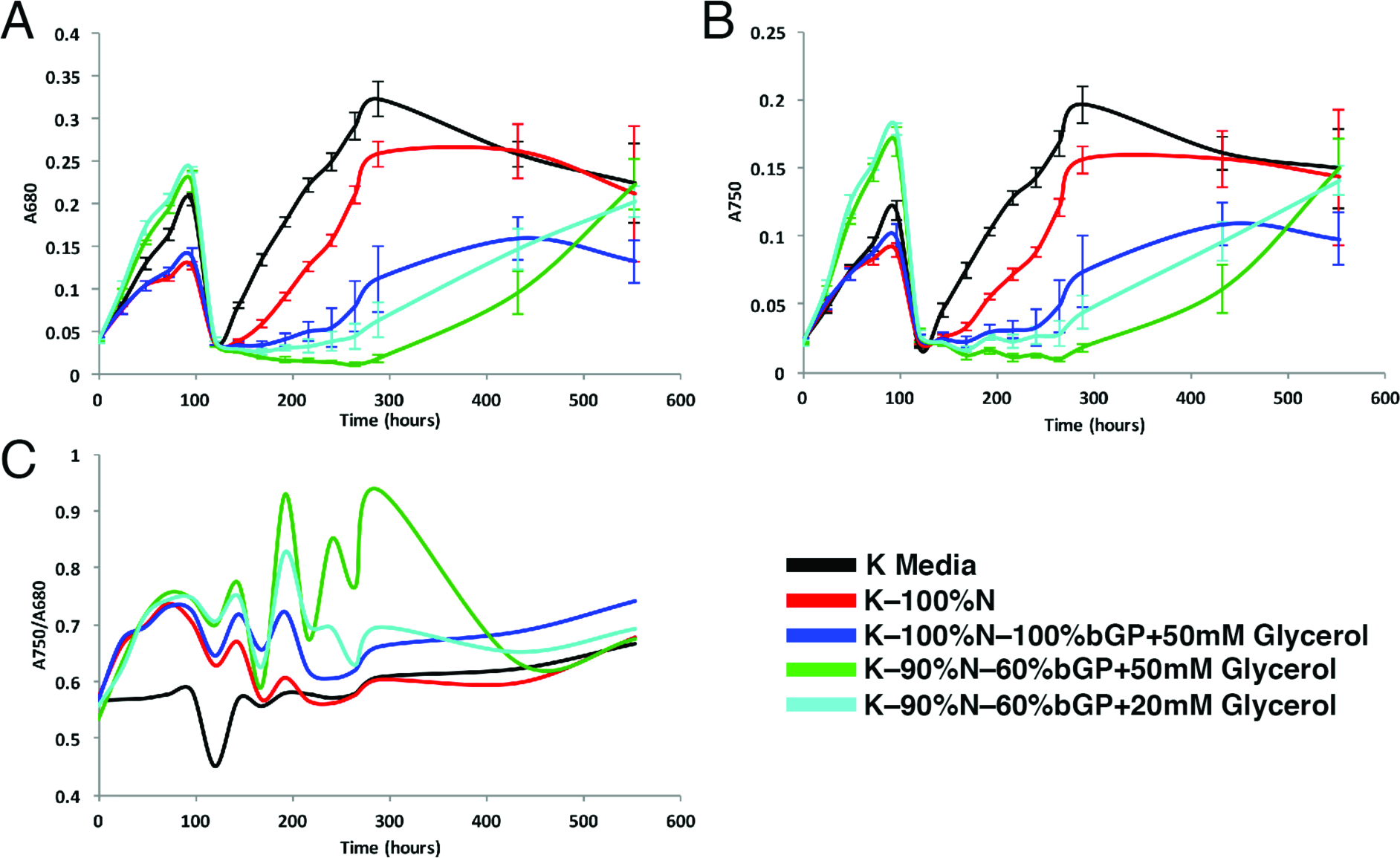
Recovery of normal growth following a batch dilution from starvation conditions. Following 120 hours of incubation in various media, each culture was spun down, washed in normal K media twice, then resuspended in fresh K media to track recovery for the next 432 hours. All initial conditions supplemented with glycerol showed a lag time of several days but then grew to standard density. Error bars represent the error in five biological replicate cultures.

**Supplemental Movie 1:** Movie of O. tauri cell caught in process of extrusion. X-ray tomogram of cell containing oriented lipid droplets in a pea-pod like structure grown in K-90%N-60%P+50mM Glycerol for 96 hours.

